# Age-associated microglia populations identified from several single cell transcriptome data

**DOI:** 10.1101/2025.07.28.665226

**Authors:** Ayaka Sugai, Hinase Moridono, Merve Bilgic, Yukiko Gotoh, Yusuke Kishi

**Affiliations:** Laboratory of Molecular Neurobiology, Institute for Quantitative Biosciences, The University of Tokyo, Tokyo 113-0032, Japan; Faculty of Pharmaceutical Sciences, The University of Tokyo, Tokyo 113-0033, Japan; Laboratory of Molecular Biology, Graduate School of Pharmaceutical Sciences, The University of Tokyo, Tokyo 113-0033, Japan; International Research Center for Neurointelligence (WPI-IRCN), The University of Tokyo, Tokyo 113-0033, Japan

**Author notes:** These authors equally contributed to this work (Ayaka Sugai, Hinase Moridono).

**Keywords:** Aging, brain, microglia, single cell transcriptome, inflammation

## Abstract

Microglial senescence contributes to inflammation and various neurodegenerative diseases. Recent single-cell transcriptome data have revealed age-associated microglial substates (AAMs) and their potential roles in the development of neurodegenerative diseases. However, the characteristics identified in each study are not necessarily consistent. Here, we perform an integrative analysis of seven previously reported single-cell RNA-seq and four single-nucleus RNA-seq datasets of microglia from young and aged mouse brains. We identify four common AAMs across all datasets and two dataset-specific AAMs. Each AAM exhibits distinct transcriptomic patterns, including alterations in ribosomal genes, *Apoe*, cytokine genes, interferon-responsive genes, and phagocytosis-related genes. Time-series and pseudotime analyses indicate that the production of AAMs is initiated by the upregulation of ribosomal genes. Predictions based on single-cell transcriptomic data of age-associated manipulations reveal an increase in specific AAMs in a stimulation-type-dependent manner. We also identify similar AAMs in human brains. Altogether, our large-scale integrative analysis highlights promising age-associated microglial populations, which may serve as novel therapeutic targets for age-related neurodegenerative diseases.

## Introduction

Brain aging is a major risk factor for several age-associated psychiatric diseases, including neurodegenerative diseases such as Alzheimer’s disease (AD) and Parkinson’s disease (PD)^1^, and the development of effective treatments for these conditions is urgently required. In the central nervous system (CNS), microglia play an essential role in maintaining brain health and function. However, with aging, microglia lose their homeostatic molecular signature and exhibit functional impairments ^2,3^, such as chronic increase in production of proinflammatory cytokines, further exacerbated by inflammatory challenges ^4–6^, elevated generation of reactive oxygen species (ROS) ^7,8^, and impaired phagocytosis ^9–11^. Therefore, a comprehensive understanding of aging-related phenotypic alterations of microglia is crucial for elucidating the mechanisms underlying brain aging and the development of age-associated neurodegenerative diseases.

Microglia are a heterogeneous cell population that varies in a spatial- and temporal-dependent manner, and recent studies have identified microglia specific to the aged brain; in this study, we call the populations as age-associated microglia (AAM). Lipid-droplet-accumulating microglia (LDAM), characterized by lipid droplet accumulation, are defective in phagocytosis, produce high levels of reactive oxygen species, and secrete proinflammatory cytokines ^8^. Recent single-cell RNA sequencing (scRNA-seq) analyses of young and aged mouse brains have also revealed AAM populations ^3^, such as those enriched in inflammatory or interferon signaling pathways have been commonly identified ^12,13^. scRNA-seq analyses of aged brains have shown that these microglial populations preferentially reside in white matter rather than gray matter ^14,15^, although two spatial transcriptome analyses have yielded differing conclusions regarding the contribution of white matter to AAMs ^16,17^. Disease-associated brains also contain specific microglial populations. For example, in the brains of AD models, such as 5xFAD or App NL-G-F transgenic mice, and in ALS model brains with *SOD1* mutation, specific microglial populations share transcriptional patterns with inflammatory AAMs rather than interferon AAMs ^13,14,18^.

In previous reports, it has not been clarified which upstream events lead to the emergence of AAM. On the other hand, inflammation is one of the most significant accelerators of brain aging ^19^. In addition, exchanging circulating factors between young and aged mice through parabiosis surgery has been shown to either reduce or improve brain function in aged or young mice, respectively ^20–22^. Acute dietary restriction or exercise has also been demonstrated to improve brain function in aged mice ^23^. However, it is still unclear which of these events function as upstream regulators for each AAM substate.

As outlined above, recent advances in single-cell analyses have revealed the characteristics of AAMs. However, several questions remain. Do AAMs identified in different studies represent the same substates, or are there additional and uncharacterized AAMs? Do different age-associated manipulations, such as Aβ accumulation and inflammation, affect microglial aging in similar or distinct ways? How do premature aging or rejuvenating interventions, such as parabiosis with young mice and dietary restriction, influence microglial aging? Addressing these questions requires a robust definition of AAMs, as microglia are a vulnerable cell type that can be affected by dissociation and isolation processes ^24^. The differences in AAMs reported across studies may stem from variations in experimental conditions, such as sex, brain regions, isolation procedures, and scRNA-seq platforms.

In this study, we performed a large-scale integrative analysis using seven scRNA-seq and four snRNA-seq datasets of young and aged mouse brains, identifying and confirming six AAM populations. This robust definition of AAMs enabled us to detect alterations in specific AAMs under various age-associated or rejuvenation interventions. Additionally, we identified similar microglial populations in the human brains of patients with neurological diseases. Therefore, our large-scale integrative dataset provides valuable insights into AAMs, contributing to a better understanding of their roles in brain aging and disease.

## Results

### Identification of AAMs by Integrative Analysis of scRNA-seq

Previous studies have identified several AAMs in mice using scRNA-seq ^3^, but their generality and functions remain unclear. To robustly examine the heterogeneity of aged microglia, we performed an integrative analysis of scRNA-seq data from young and aged mice using seven independent public datasets ^12,18,25–29^ (Fig. 1A, Supplementary Table 1). These datasets include microglia from two-to six-month-old young and 18-to 29-month-old aged wild-type C57BL/6 mice. However, the datasets differ in terms of gender, brain regions (whole brain, hippocampus, or subventricular zone), and experimental conditions, such as dissociation methods (enzyme-based or direct homogenization), cell isolation methods (with or without microglial sorting), and scRNA-seq platforms (10x Chromium-based drop-seq or MARS-seq) (Supplementary Table 1). Microglial data were extracted from each dataset and integrated using the same number of cells (1,769 cells per dataset) with Seurat software ^30^, and we identified 15 clusters in the integrated dataset (Fig. 1B, Extended Data Fig. 1A)^30^. The integrated data revealed distinct distributions of young and aged microglia in UMAP space (Fig. 1C). The distributions of individual datasets overlapped significantly (Extended Data Fig. 1B), indicating successful integration.

**Figure 1.**
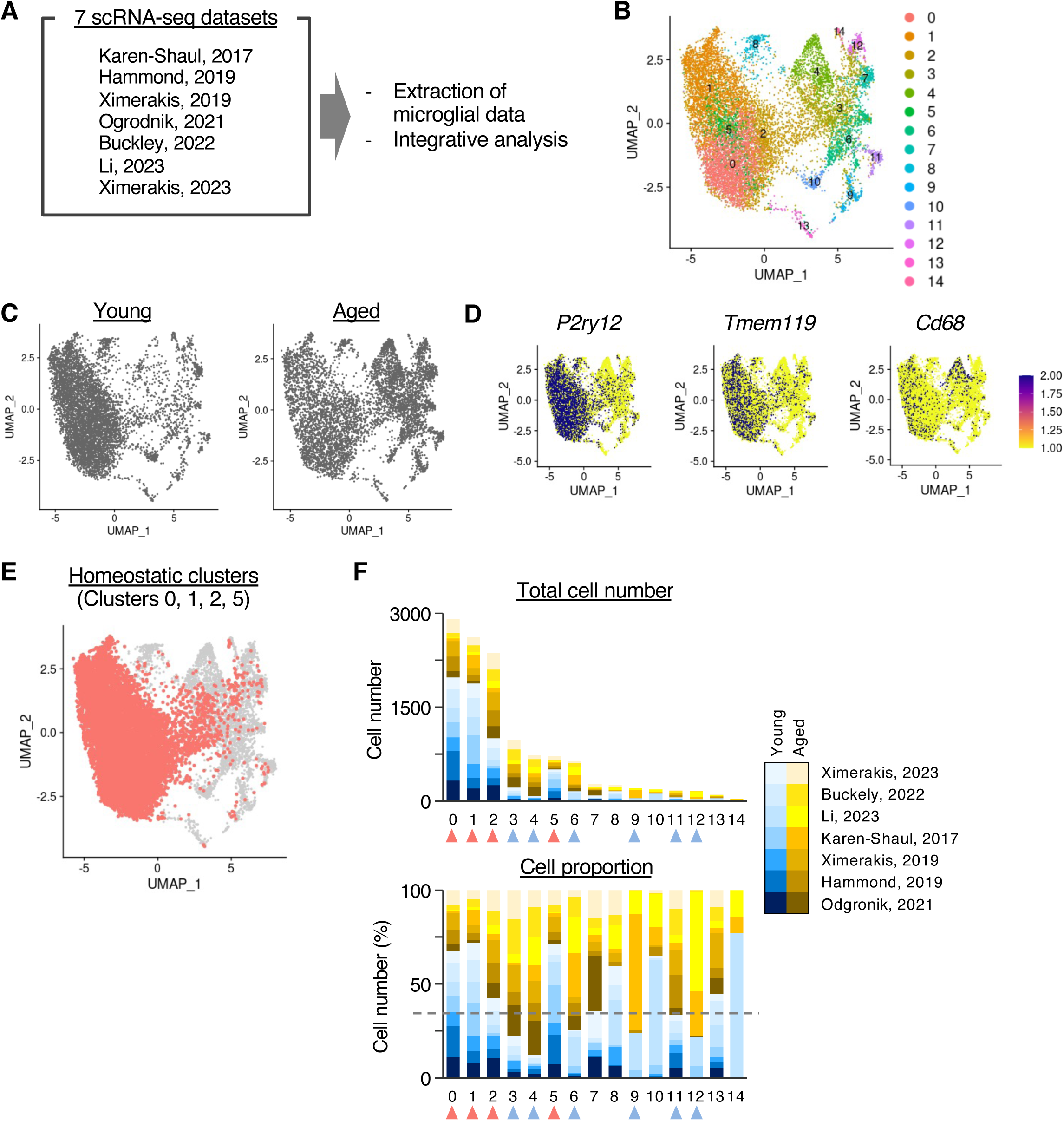
Integration of seven public scRNA-seq datasets of young and aged microglia. **A.** Schematic representation of the integration of seven public scRNA-seq datasets. **B.** Visualization of clusters in UMAP space. **C.** Distribution of young and aged microglia in UMAP space. Dark gray dots indicate young or aged microglia in the left or right panels, respectively. **D.** Expression patterns of homeostatic (*P2ry12* and *Tmem119*) and activated (*Cd68*) microglia markers. **E.** Distribution of homeostatic microglia (clusters 0, 1, 2, and 5) in UMAP space. **F.** Total cell numbers (upper panel) and percentages (lower panel) of cells in each cluster. Different colors indicate distinct ages and datasets. Red or light blue arrowheads represent homeostatic microglia or AAMs, respectively.

Clustering analysis based on gene expression patterns categorized microglia into 15 clusters (Fig. 1B). The expression levels of *P2ry12* and *Tmem119*, markers for homeostatic microglia, and *Cd68*, a marker for activated microglia, showed that clusters 0, 1, 2, and 5 were homeostatic microglia clusters enriched with young microglia (Fig. 1D, E). Clusters with approximately twice as many microglia in aged brains as in young brains were defined as AAM clusters, specifically clusters 3, 4, 6, 9, 11, and 12 (Fig. 1F). Among these, clusters 9 and 12 were predominantly derived from two datasets of Keren-Shaul et al. and Li et al. (Fig. 1F), suggesting they may appear under specific conditions. We checked the difference in mouse and experimental conditions (Supplementary Table 1), however, could not find common difference between Shaul et al. and Li et al. and other datasets, and the conditions under which cluster 9 and cluster 12 appear or can be detected are unknown at present. In contrast, clusters 3, 4, 6, and 11 included aged microglia from all datasets, indicating their commonality in aged brains under diverse experimental conditions (Fig. 1F, Extended Data Fig. 1C). Collectively, the integrative analysis revealed four common AAMs (clusters 3, 4, 6, and 11) and two context-dependent AAMs (clusters 9 and 12).

### Different Properties of AAMs

We determined the properties of each AAM by analyzing differentially expressed genes (DEGs) using the Seurat package, comparing homeostatic microglia (clusters 0, 1, 2, and 5) and each AAM (clusters 3, 4, 6, 9, 11, or 12) (Fig. 1D-E; Fig. 2B, C, E, G; Extended Data Fig. 2E, H; Supplementary Table 2). Four out of six AAMs (clusters 3, 4, 6, and 12) showed high expression of ribosomal genes, such as *Rpl12*, *Rpl38*, and *Rps2* (Extended Data Fig. 2A). While homeostatic microglia also expressed ribosomal genes, their expression levels in these AAMs were higher. Gene ontology (GO) analysis confirmed that upregulated genes in clusters 3, 4, 6, and 12 were enriched for terms such as “Cytoplasmic Translation” and “Peptide Biosynthetic Process” (Fig. 2A, Supplementary Table 3). This is consistent with findings from bulk RNA-seq analyses, which also report ribosomal gene upregulation ^31^ and higher translational activity in aged microglia ^32^. As these four AAMs accounted for 88% of the total AAM population, we propose that increased ribosomal gene expression is the most common feature of aging-related phenotypic alterations of microglia. We designate these four AAMs as Type1 AAMs.

**Figure 2.**
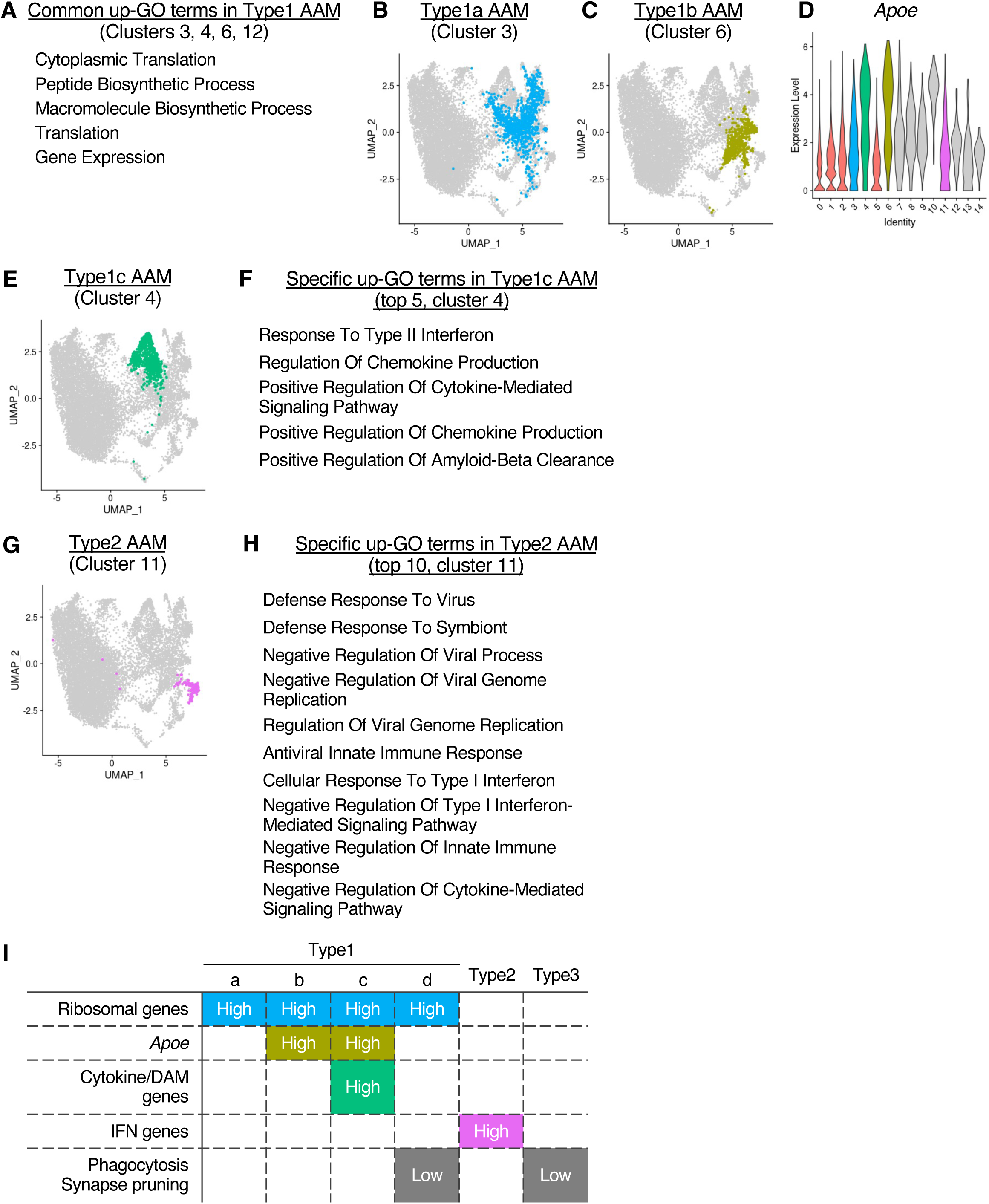
Characteristics of common AAMs. **A–D**. Distribution of the four common AAMs in UMAP space. **E.** Top five Biological Process GO terms of upregulated DEGs among Type1 AAMs. **F.** Expression levels of the Apoe gene in each cluster. **G.** Top five Biological Process GO terms of upregulated DEGs in Type1c AAMs. **H.** Top ten Biological Process GO terms of upregulated DEGs in Type2 AAMs. **I.** Summary of the characteristics of DEGs in each AAM.

Among the Type1 AAMs, transcriptional differences were evident. Cluster 3 exhibited no notable characteristics other than ribosomal gene upregulation. In contrast, clusters 4 and 6 displayed high expression of *Apoe* (Fig. 2D), a gene that is highly expressed in aged and AD microglia, and its expression has an important role in AD-related abnormal behavior ^33^. Except for the characteristic of higher *Apoe* expression, the DEGs in cluster 6 overlapped with those in cluster 3 (28 out of 34 up-DEGs), suggesting similar characteristics of higher ribosomal genes (Supplementary Tables 2, 3). However, cluster 4 also expressed cytokine-related genes, as confirmed by GO analysis (Fig. 2F, Extended Data Fig. 2B). This is similar characteristics with previously identified AAMs such as disease-associated microglia (DAM), aged cluster 2 (OA2), microglial neurodegenerative phenotype (MGnD), and white matter-associated microglia (WAM) ^12,14,18,34^. Key DEGs in cluster 4, including *Csf1*, *Cst7*, *Itgax*, and *Lpl*, were also highly expressed in these previously reported AAMs (Extended Data Fig. 2B). We confirmed the increase of cluster 4 microglia during brain aging by staining Lpl protein, which was enriched in the cluster 4, using young (9-10 weeks old) and aged (112-116 weeks old). Lpl-and Iba1-positive microglia significantly increased in dentate gyrus of aged brains and showed increasing tendency in CA1/CA3, in addition, Lpl-negative and Iba1-positive microglia also increased in aged brains (Extended Data Fig. 2C, D).

Cluster 12, another Type1 AAM, did not exhibit high expression of *Apoe* or cytokine genes. Instead, its downregulated DEGs were associated with phagocytosis and synapse pruning (Extended Data Fig. 2F). Impaired phagocytosis and synapse pruning are hallmark features of aged microglia ^2^. Based on these findings, we designate clusters 3, 6, 4, and 12 as Type1a, 1b, 1c, and 1d AAMs, respectively.

Other AAMs, cluster 9 and 11, did not highly express ribosomal genes, and cluster 11 highly expressed innate immune system-related genes as shown in the enriched GO terms, such as “Defense Response To Virus” and “Cellular Response To Type I Interferon” (Fig. 2H). The population with higher expression of innate immune system- and interferon-related genes is also observed in the previous reports, called as OA3 and interferon response microglia (IRM) ^12,13,15^. The upregulated DEGs in cluster 11, such as *Ifit2*, *Ifit3*, *Irf7*, and *Oasl2*, were also identified as DEGs in OA3 and IRM (Extended Data Fig. 2G). We designate cluster 11 as Type2 AAM.

The downregulated DEGs of cluster 9 were enriched with genes related to synapse pruning (Extended Data Fig. 2I). We designate cluster 9 as Type3 AAM. Both clusters 9 and 12 exhibited lower expression of phagocytosis and synapse pruning genes and were observed in two out of seven datasets. Previous studies have reported that microglia isolated from aged brains exhibit reduced phagocytic activity for Aβ peptides in in vitro culture compared to those from young brain ^9,10^. However, this finding is still controversial due to the challenges in accurately evaluating phagocytic activity in vitro^11^. Our results suggest that a decline in phagocytic activity is not universally observed in aged mice but appears under specific conditions.

Focusing on the proportions of each AAM across datasets, Type1c was consistently present at a certain proportion (6.5–16.9%) in all seven datasets (Extended Data Fig. 1C). In contrast, the proportions of Type1a and Type1b displayed greater variability between datasets, ranging from 1.5% to 22.4% for Type1a and from 1.9% to 19.0% for Type1b. However, the combined proportion of Type1a and Type1b exceeded 10% in all datasets. This indicates that *Apoe* expression depends on the specific conditions of the aged mice, which in turn determines whether Type1a or Type1b becomes dominant, however, we could not find the difference in mouse and experimental conditions between Type1a- or Type1b-enriched datasets (Supplementary Table 1). Type2 AAMs had the lowest proportions, with values ranging from 0.9% to 3.8%, and also showed variation across datasets. Notably, all AAMs were observed in young mice, indicating that AAMs are not exclusive to aged microglia and may appear at earlier stages of development.

In summary, we identified six distinct AAM populations with varying properties characterized by differences in the expression levels of ribosomal genes, *Apoe*, cytokine genes, interferon-related genes, phagocytosis genes, and synapse pruning genes (Fig. 2I).

### AAMs in snRNA-seq Datasets

Recently, snRNA-seq has become the preferred method for neuroscience research, as it avoids the challenges of recovering neuronal cell bodies during dissociation. To assess AAMs in snRNA-seq datasets, we integrated four public datasets from young (1–4 months old) and aged (18–24 months old) mice, resulting in the identification of 11 clusters ^16,17,25,35^ (Fig. 3A-B). We designated snRNA-seq clusters as n0, n1, etc., to distinguish them from scRNA-seq clusters. The distributions of young and aged microglia were largely similar, with slight differences (Fig. 3C). Based on the expression level of *P2ry12*, a homeostatic microglia gene, we identified clusters n0, n1, and n2 as homeostatic microglia (Fig. 3D, E). Among the clusters, only n6 contained more than 65% aged microglia and was present across all four datasets (Fig. 3F, G).

**Figure 3.**
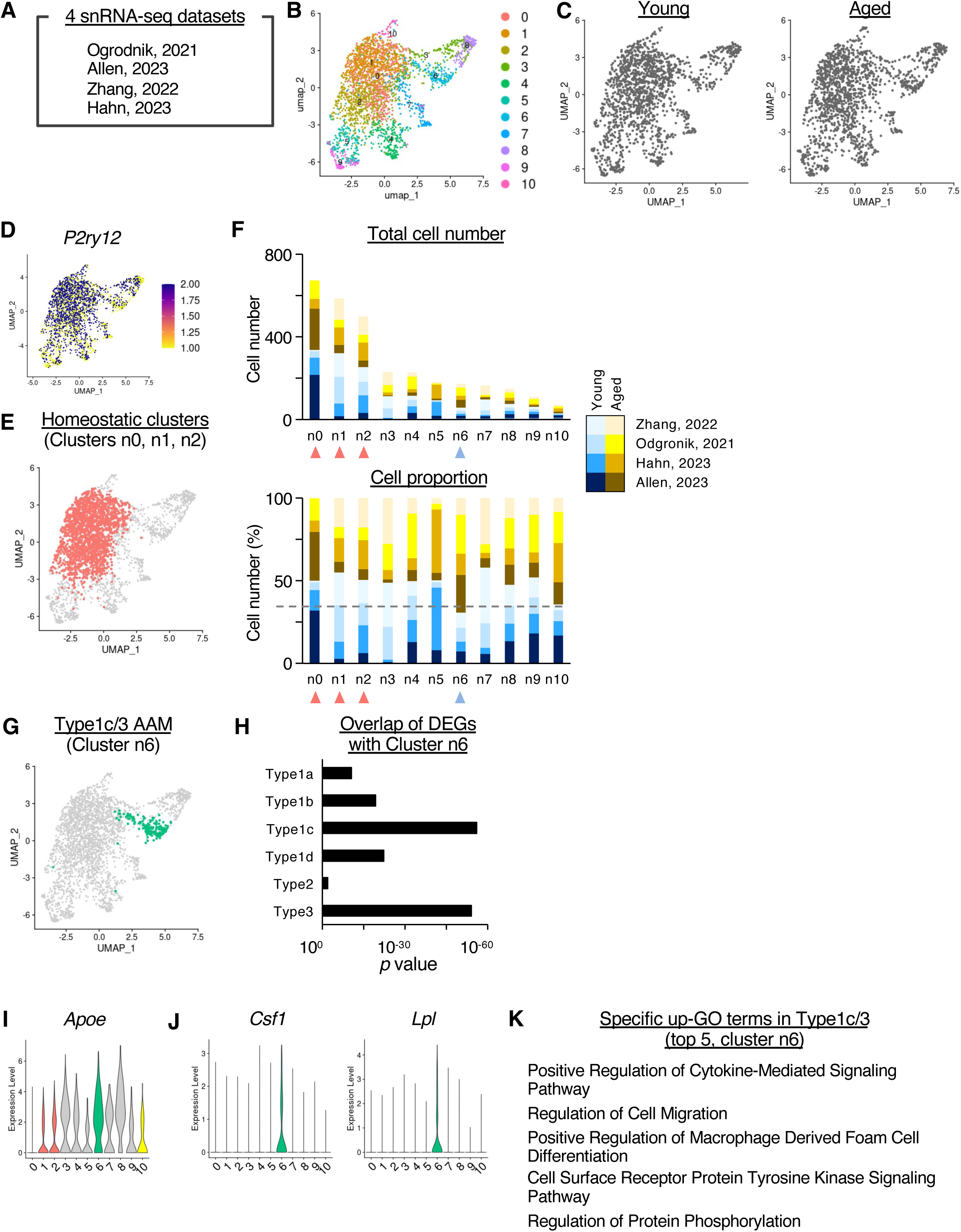
Integration of four public snRNA-seq datasets of young and aged microglia. **A.** Schematic representation of the integration of four public snRNA-seq datasets. **B.** Visualization of clusters in UMAP space. **C.** Distribution of young and aged microglia in UMAP space. Dark gray dots indicate young or aged microglia in the left or right panels, respectively. **D.** Expression patterns of *P2ry12*, a marker for homeostatic microglia. **E.** Distribution of homeostatic microglia (clusters n0, n1, and n2) in UMAP space. **F.** Total cell numbers (upper panel) and percentages (lower panel) of cells in each cluster. Different colors indicate distinct ages and datasets. Red or light blue arrowheads represent homeostatic microglia and AAMs, respectively. **G.** Distribution of cluster n6, an AAM identified by snRNA-seq, in UMAP space. **H.** Significance of the overlap between the DEGs of cluster n6 and the six AAMs identified by scRNA-seq.

Comparative analysis revealed that DEGs in cluster n6 overlapped most with those in Type1c and Type3 AAMs from scRNA-seq (Fig. 3H, Supplementary Table 4), suggesting that cluster n6 is a mixture of Type1c and Type3 AAMs. We designate cluster n6 as Type1c/3 AAM. Cluster n6 exhibited similar properties to Type1c, including high expression of *Apoe*, *Csf1*, and *Lpl*, and enrichment of cytokine genes (Fig. 3I-K, Supplementary Table 5). However, ribosomal and interferon-related genes were not detected in any snRNA-seq clusters (Extended Data Fig. 3B, C). This limitation aligns with findings that snRNA-seq cannot fully capture microglial activation states ^36^. These results suggest that snRNA-seq datasets are useful for analyzing Type1c and Type3 AAMs but are inadequate for identifying all AAMs detected by scRNA-seq.

### Lineage Relationship Between AAMs

We next investigated the lineage relationships among AAMs. First, we analyzed the timing of the emergence of each AAM using two time-series scRNA-seq datasets ^28,29^. Buckley et al. provided scRNA-seq data in subventricular zone from 28 mice, tiling 26 different ages from 3 months to 29 months, and we categorized them into five stages: 3-5, 6-10, 12-16, 18-22, and 23-29 months old. From Li et al., we used scRNA-seq data in whole brain of PBS-injected control mouse at 3, 14, and 25 months old. In both datasets, Type1c AAMs emerged after 24 months, whereas Type1a and Type1b AAMs were observed to increase prior to the emergence of Type1c AAMs, at 18-22 months old in Buckley data or 14 months old in Li data (Fig. 4A, B). We further conducted pseudotime analysis using our integrated data (Fig. 4C), which revealed that the first node, indicated with yellow arrow, from the homeostatic population was located in the Type1a cluster. The lineage then branched into two trajectories: one leading to Type1b, Type2, and Type3, and the other leading to Type1c and Type1d. Both the time-series and pseudotime analyses indicate that during brain aging, Type1a AAMs emerge first and subsequently differentiate into other AAMs with distinct properties. Previous studies have shown that increased translational activity induced by the mTOR pathway contributes to the aberrant expression of cytokine proteins ^32^. Thus, the emergence of Type1a AAMs, driven by increased ribosomal gene expression, may provide the foundation for the higher expression of AAM-related proteins, such as cytokines in Type1c AAMs.

**Figure 4.**
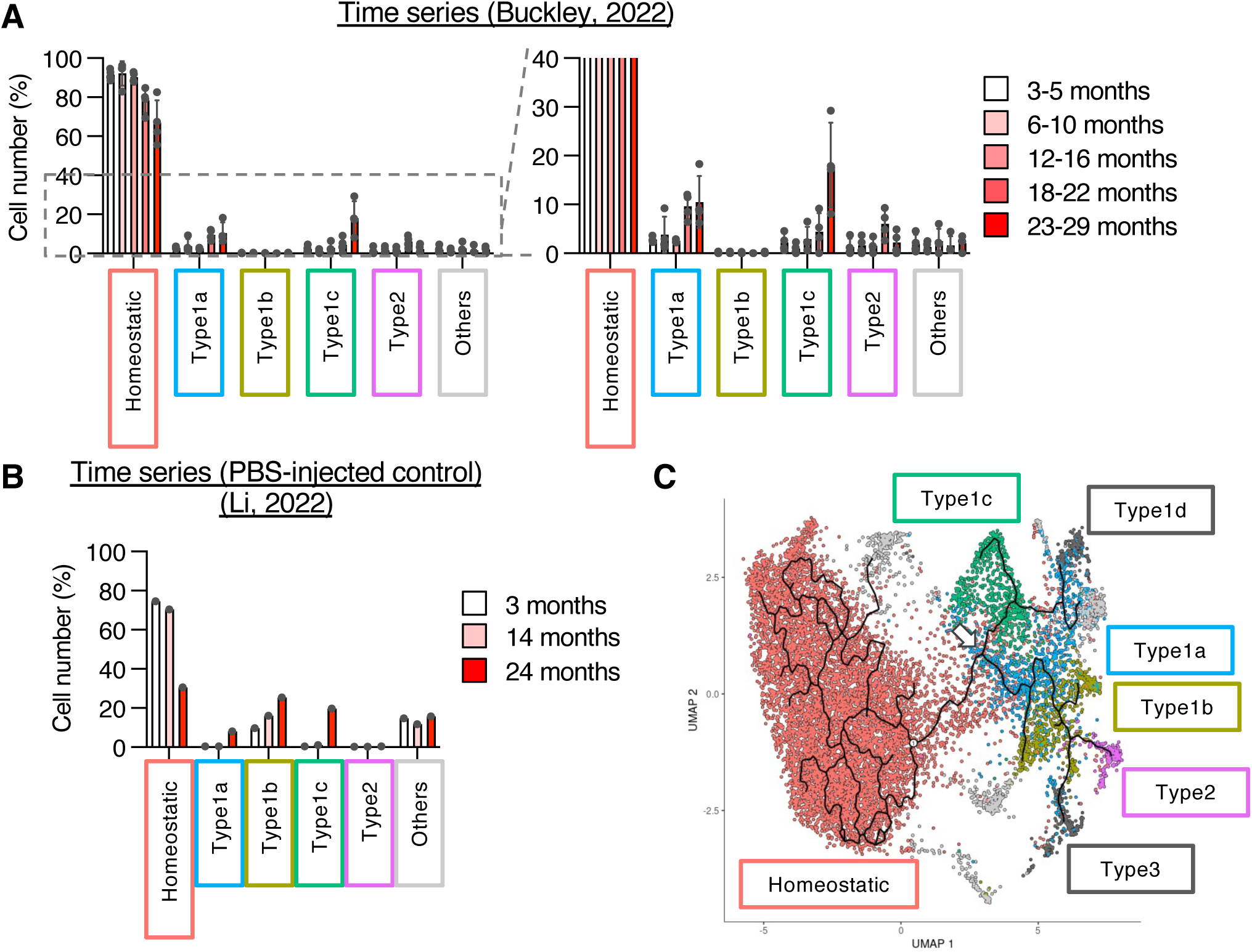
Emergence of AAMs during aging and pseudotime analysis of young and aged microglia. **A–B**. Prediction of corresponding clusters (homeostatic microglia, four common AAMs, and others) for cells from scRNA-seq performed at multiple time points spanning young and aged stages. Bar graphs show the number of predicted cells. Data are presented as mean ± SD. **C**. Pseudotime analysis of young and aged microglia integrated in Figure 1. Black circles represent nodes, and black lines indicate predicted lineage trajectories. The white arrow indicates the nodes firstly branched in age-associated clusters.

### The Effect of Age-Associated and Rejuvenation Manipulations on AAMs

We next examined the factors driving the emergence of each AAM population. Since aging-related phonotype can be modulated by various interventions, we analyzed the effects of age-associated manipulations on the homeostatic microglia and four common AAMs, Type1a, 1b, 1c, and Type2 (Fig. 5A).

**Figure 5.**
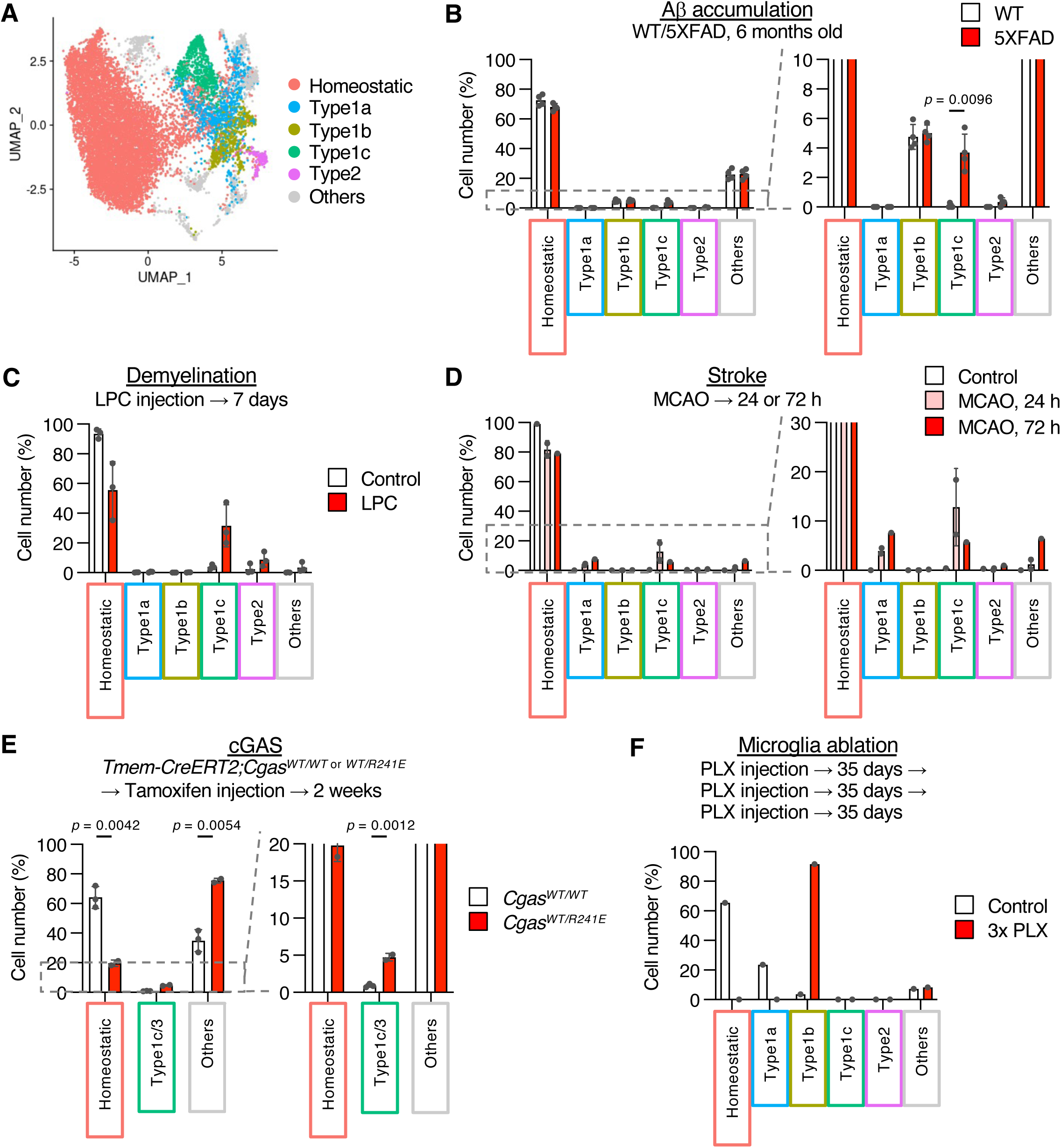
The effect of age-associated manipulations on the number of homeostatic microglia and four common AAMs. **A.** Distribution of homeostatic microglia and the four common AAMs in UMAP space. **B–F**. Prediction of corresponding clusters (homeostatic microglia, four common AAMs, and others) for cells from scRNA-seq of 5XFAD AD model mice (B), LPC injection-demyelination model mice (C), MCAO-stroke model mice (D), mice expressing constitutively active cGAS (E), and faster microglia turnover model mice (F). Bar graphs show the number of predicted cells. Data are presented as mean ± SD. *p*-values were calculated using multiple unpaired t-tests with Welch correction.

Accumulation of Aβ is known to accelerate brain function decline ^37^. Using scRNA-seq datasets from Aβ accumulation model mice (5XFAD mice), we classified the cells into homeostatic microglia and four common AAMs. Specifically, we performed label transfer in Seurat based on prediction scores and determined the difference of the number of microglia predicted as homeostatic microglia or four common AAMs. ^18,30,38^ (Fig. 5B). As previously reported, the number of Type1c AAMs, which share properties with DAM, significantly increased, whereas no significant changes were observed in Type1a, Type1b, or Type2 AAMs.

Chronic inflammation is a hallmark of the aged brain ^19^. We analyzed demyelination and stroke model mice, which induce acute brain inflammation ^12,39^. Type1c AAMs were significantly increased in both models: demyelination induced by bilateral injection of lysolecithin (LPC) into subcortical white matter and stroke induced by middle cerebral artery occlusion (MCAO) (Fig. 5C, D). Type2 AAMs increased in response to LPC injection, while Type1a AAMs increased in response to MCAO (Fig. 5C, D). Furthermore, inflammation induced by the expression of a constitutively active form of cGAS also increased Type1c/3 AAMs in snRNA-seq analysis (Fig. 5E) ^40^. These results suggest that Aβ accumulation and inflammation are associated with the emergence of Type1a, Type1c, and Type2 AAMs.

Acceleration of microglial turnover by repetitive microglial ablation further enhanced brain aging ^28^. Ablation performed three times using PLX5622 with 35-day intervals significantly increased the number of Type1b AAMs while decreasing the number of Type1a AAMs ^28^ (Fig. 5F).

Rejuvenation strategies, such as sharing circulating factors with young mice, exercise, and dietary restriction (DR), can partially restore brain function in aged mice^23^. We analyzed the effects of heterochronic parabiosis between young and aged mice compared to isochronic parabiosis using three scRNA-seq datasets from two groups ^27,29^. In one dataset, aged mice fused with young mice showed no significant changes in the numbers of the four common AAMs (Fig. 6A). In another dataset, Type1a AAMs decreased (Fig. 6B), while in a third dataset, both Type1c and Type2 AAMs decreased (Fig. 6C). These results suggest that the effects of heterochronic parabiosis on AAM numbers may vary depending on experimental conditions. Exercise did not alter the numbers of the common AAMs ^29^ (Extended Data Fig. 4A). In contrast, snRNA-seq analysis of brains subjected to acute and long-term DR showed a decrease in Type1c/3 AAMs ^17^ (Fig. 6D).

**Figure 6.**
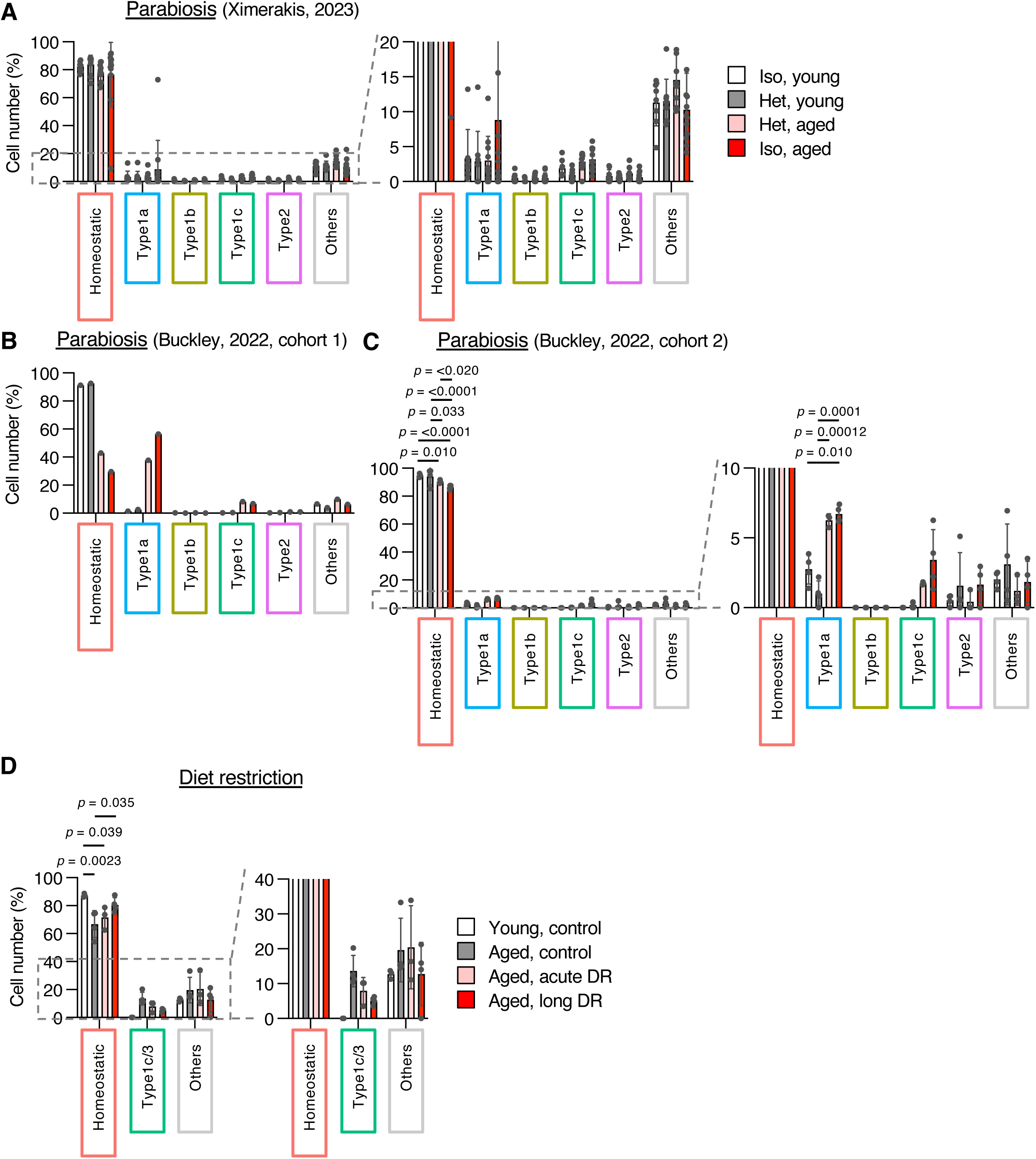
The effect of rejuvenation and premature aging manipulations on the number of homeostatic microglia and four common AAMs. **A–D**. Prediction of corresponding clusters (homeostatic microglia, four common AAMs, and others) for cells from scRNA-seq of parabiosis between young and aged mice (A–C) and DR mice (D). Bar graphs show the number of predicted cells. Data are presented as mean ± SD. Adjusted *p*-values were calculated using two-way ANOVA followed by Tukey’s multiple comparison test.

In summary, specific AAM populations responded differently to various manipulations. Parabiosis affected Type1a and Type1c AAMs depending on experimental conditions, while DR consistently reduced Type1c/3 AAMs. Using public scRNA-seq and snRNA-seq datasets, we identified upstream pathways regulating AAM emergence: Aβ accumulation and inflammation increased Type1c AAMs, accelerated microglial turnover increased Type1b AAMs, and DR reduced Type1c AAMs. These findings highlight the utility of our integrated single-cell data in determining the roles of brain manipulations on AAMs.

### AAMs in Human Neurological Diseases

Finally, we investigated whether AAMs identified in this study exist in the human brain. Corresponding clusters in our mouse integrated dataset were compared to microglia in human scRNA-seq data obtained from the dorsolateral prefrontal cortex of patients with mild cognitive impairment (MCI) and Alzheimer’s disease (AD). Homologous genes between the two species were used to identify matching clusters ^41^ (Extended Data Fig. 5A). Consistent with previous studies ^18,41^, we observed a higher abundance of Type1c AAMs, corresponding to DAM, in AD patients compared to MCI patients (Extended Data Fig. 5A). Furthermore, mouse AAMs identified in our study were also found in several other human neurological diseases using additional datasets^42^ (Extended Data Fig. 5B). These results demonstrate the potential applicability of our mouse AAM dataset to human datasets for analyzing the impact of neurological diseases on AAMs.

## Discussion

Microglia play a central role in brain aging, and previous studies have identified AAM substates individually. However, no unified analysis of AAMs has been conducted, and discussions regarding the relationships among AAMs are lacking. One pioneering study performed an integrative analysis of mouse scRNA-seq datasets spanning several developmental stages, from embryonic to aged stages, as well as neurodegenerative model mice, and identified previously reported DAM populations composed of two distinct substates, referred to as DAM and disease inflammatory macrophage (DIM) in their study ^43^. However, their focus on development, aging, and disease included only one aged dataset. In contrast, the present study integrated seven scRNA-seq datasets and identified four common AAMs across all datasets. Among these, Type1c and Type2 AAMs correspond to previously reported populations, such as DAM, OA2, MGnD, or WAM, and OA3 or IRM ^12–15,18,34^, respectively. However, Type1a and Type1b AAMs represent novel substates identified in this study.

Prior research has demonstrated the role of mTOR-mediated translational activity in aged microglia in relation to brain aging phenotypes, such as altered locomotor activity ^32^. Considering our findings of elevated ribosomal gene expression in Type1 AAMs and the positioning of Type1a as an upstream precursor to other AAMs, as suggested by time-series and pseudotime analyses, we propose that upregulation of ribosomal genes and translational activity act as key drivers of aging-related phenotypic alterations of microglia. Indeed, the mTOR pathway, a regulator of translational activity, is considered a critical factor in aging. Treatment with rapamycin, an mTOR inhibitor, has been shown to extend lifespan in mice and ameliorate neurodegeneration-associated neuronal death and cognitive deficits ^44–47^. This raises the question: what age-associated environmental factors induce ribosomal gene upregulation? Based on our prediction analyses of age-associated manipulation datasets, factors such as stroke, accelerated microglial turnover, and aging-associated changes in circulating factors may upregulate Type1a or Type1b AAMs, potentially driving ribosomal gene expression and increased translational activity.

Our analysis revealed higher *Apoe* expression levels in Type1b and Type1c AAMs. While the upregulation of *Apoe* in Type1c is consistent with previous reports, the primary distinction between Type1a and Type1b lies in the elevated *Apoe* expression in Type1b. Despite comparable total cell numbers of Type1a and Type1b across datasets, their relative proportions varied, suggesting that *Apoe* expression is influenced by breeding and experimental conditions. *Apoe* expression in microglia is known to be upregulated by stress, injury, Aβ plaque accumulation, and *Apoe* expression in other cell types, such as neurons ^33^. The precise mechanisms triggering *Apoe* upregulation in addition to ribosomal gene activation during brain aging remain unknown and warrant further investigation. While we observed an increase in Type1c AAMs in AD compared to MCI, the presence of Type1a AAMs was limited, and Type1b AAMs were more prevalent in human datasets (Extended Data Fig. 5A, B). However, these observations are challenging to generalize, as the majority of the human data analyzed were derived from aged patients (e.g., all patients in Olah, 2020, were over 79 years old, and 84% of patients in Tuddenham, 2024, were over 61 years old). This suggests that the regulatory mechanisms of *Apoe* expression may differ between mouse and human microglia.

In summary, our integrative analysis of young and aged microglia identified four common and two context-specific AAM substates, highlighting the importance of the newly identified Type1a substates in the emergence of AAMs. Additionally, our dataset provides a valuable platform for investigating the effects of brain manipulations on AAMs. Although our conclusions regarding the functions and lineage relationships of AAMs are primarily based on scRNA-seq data, in vivo manipulation of AAMs in aged brains remains challenging. Future studies are needed to elucidate the roles of AAMs identified in this study.

## Materials and Methods

### Acquiring Public scRNA-seq and snRNA-seq Datasets, Quality Check, Clustering, and Visualization in UMAP Space

The datasets of scRNA-seq and snRNA-seq used in this study are listed in Supplementary Table 1. Data processing and visualization were performed using the Seurat package (version 4.3.0) in R (version 4.2.0) ^30^. After obtaining original cell-gene matrices from the database, quality control was conducted in Seurat, retaining genes expressed in at least three cells per dataset and cells with 200 to 5,000 detected genes and fewer than 20% mitochondrial genes. Total unique molecular identifiers (UMIs) were normalized to 10,000 using the ‘LogNormalizè function, and expression levels were standardized to a mean of 0 and a standard deviation of 1 using the ‘ScaleDatà function. Highly variable genes (2,000 per dataset) were identified using the ‘FindVariableFeatures‘ function. After dimensionality reduction via the ‘RunPCÀ function, principal components (PCs) were selected for clustering using the ‘FindNeighbors‘ and ‘FindClusters‘ functions. The clustering conditions, such as resolutions and used dimentions, were dependent on each dataset and shown in Supplementary Table 1. Other used settings were default. Then, the results were visualized in UMAP space using the ‘RunUMAP‘ function.

### Extraction of Microglial Data

Microglial data were extracted using the ‘subset‘ function in Seurat. Clusters expressing *Tmem119* and *Aif1*, markers for microglia, were retained. To exclude monocytes and macrophages, *Hexb*, a marker specific to microglia in the CNS, was also used, as it is not expressed in microglia ^48^.

### Integration of Multiple Datasets

To ensure uniform representation across datasets, 1,769 cells or 752 nuclei were randomly sampled, to balance the sample size across datasets, before integration by “sample” function from each scRNA-seq or snRNA-seq dataset, respectively. Following a previously reported “anchor-based”integration method with Seurat ^30^, genes with similar expression levels across datasets were identified using the ‘SelectIntegrationFeatures‘ function, and “anchor” cell pairs, assumed to have similar transcriptomes across datasets, were identified using the ‘FindIntegrationAnchors‘ function. The datasets were then integrated into a single object using the ‘IntegrateDatà function. Expression levels were standardized to a mean of 0 and a standard deviation of 1 using the ‘ScaleDatà function. After dimensionality reduction via the ‘RunPCÀ function, principal components (PCs) were selected for clustering using the ‘FindNeighbors‘ and ‘FindClusters‘ functions. The clustering resolution was set at 0.8. In ‘FindNeighbors‘, we used 1:30 dimensions in integrated scRNA-seq and 1:10 dimensions in integrated snRNA-seq data. Other used settings were default. Then, the results were visualized in UMAP space using the ‘RunUMAP‘ function.

### Identifying Marker Genes consistently found across datasets and GO Analysis

Common marker genes between clusters of interest were identified by the ‘FindConservedMarkers‘ function using Wilcoxon rank sum test, with a max adjusted *p*-value threshold of 0.05. Gene ontology (GO) analysis for upregulated or downregulated marker genes was performed using the Enrichr software ^49^, employing the “GO Biological Process 2023” database for scRNA-seq and the “GO Biological Process 2025” database for snRNA-seq. Top five or ten GO terms with lowest adjusted *p*-values were shown in the figures.

### Prediction of Mouse Datasets on Clusters in Integrated Data

Each data set was normalized using ‘SCTransform‘. After dimensionality reduction via the ‘RunPCÀ function, principal components (PCs) were selected for clustering using the ‘FindNeighbors‘ and ‘FindClusters‘ functions. The clustering resolution was set at 0.8, and used dimensions for each datasets are shown on Supplementary Table1. Anchor cell pairs were identified using the ‘FindTransferAnchors‘ function with SCT normalization method and the dimentions used was the same with the references. Datasets in Figures 4, 5, and 6 were transferred into young and aged integrated datasets using the ‘TransferDatà function. The dimentions used were the same with those we used in ‘FindNeighbors‘ function. Each cell was annotated into the cluster with the highest prediction score within the integrated data clusters.

### Prediction of Human Datasets on Clusters in Integrated Data

Filtered feature-barcode matrices (filtered_feature_bc_matrix) from Tuddenham, 2024, were directly downloaded, whereas those from Olah, 2020, were generated using the ‘count‘ function in the CellRanger software from raw FASTQ files. These matrices were imported into R using the ‘Read10X‘ function. Gene names from human (GRCh38.p13) and mouse (GRCm39) datasets were converted into Ensembl IDs using the ‘useMart‘ function in the biomaRt software, and corresponding gene sets were determined using the ‘getLDS‘ function. Only human genes with corresponding mouse genes were used. Converted human gene names were mapped to mouse gene names. After converting the datasets into Seurat objects, they were merged using the ‘mergè function, and predictions were performed in the same manner as for mouse datasets.

### Comparison of scRNA-seq and snRNA-seq Clusters

The overlaps between DEGs specific to AAMs in scRNA-seq and snRNA-seq clusters were analyzed using Fisher’s exact test, implemented in R with the ‘fisher.test‘ function.

### Single-cell trajectory Analysis of Integrated scRNA-seq

An integrated scRNA Seurat object was converted into a CellDataSet (CDS) object using the “new_cell_data_set” function in the Monocle3 package. Gene annotations and cell metadata were derived from the PCA loadings and count matrix dimensions, respectively. UMAP coordinates, PCA gene loadings, and cluster identities obtained from Seurat were transferred to the CDS to retain consistent dimensionality reduction and clustering results. To establish a unified trajectory, all cells were assigned to a single partition. Trajectories and pseudotime ordering were inferred using Monocle3’s “learn_graph” and “order_cells” functions. Finally, cell states were visualized within the UMAP space.

### Ethics Statement

All animal experiments were approved by the Animal Care and Use Committee of The University of Tokyo (approval numbers: P25-8 and P30-4 in the Graduate School of Pharmaceutical Sciences, and 0421 in the Institute for Quantitative Biosciences). All procedures adhered to The University of Tokyo’s guidelines for laboratory animal care and the ARRIVE guidelines.

### Mouse Maintenance

Mice were housed in a temperature- and humidity-controlled environment (23 ± 3 °C and 50 ± 15%, respectively) under a 12-hour light/dark cycle. Sterile cages (Innocage, Innovive) containing bedding chips (PALSOFT, Oriental Yeast) were used, with two to six mice per cage. Irradiated food (CE-2, CLEA Japan) and filtered water were provided ad libitum.

### Immunohistofluorescence Analysis

Immunofluorescence staining was performed as described previously ^50^. Two sets of 9- or 112-week-old C57BL/6J male mice and one set of 10- or 116-week-old C57BL/6N female mice were used. Brains were dissected, fixed in 4% paraformaldehyde (PFA) in phosphate-buffered saline (PBS) at 4 °C for over 16 hours, incubated with 15% and 30% sucrose in PBS, and embedded in O.C.T. compound (SAKURA Finetek; 4583) at -80 °C. Coronal sections (12 μm) were prepared using a cryostat (Thermo Fisher Scientific or SAKURA Finetek). Sections underwent antigen retrieval in HistoVT One (Nacalai Tesque; 06380-05) at 105 °C for 10 minutes. Samples were blocked with Tris-buffered saline (TBS) containing 0.1% Triton X-100 and 3% bovine serum albumin for 2 hours at room temperature, followed by incubation with primary antibodies at 4 °C for over 16 hours. Alexa Fluor-conjugated secondary antibodies (1:1,000, Thermo Fisher Scientific; A32766 and A32794) were applied at 4 °C for over 2 hours. Nuclei were counterstained with Hoechst 33342 (1:10,000 dilution, Thermo Fisher Scientific; H3570) before mounting in Mowiol (Calbiochem or Vector Laboratories; H-1700). Primary antibodies included anti-Iba1 (WAKO; 019-19741; 1:500 dilution) and anti-Lpl (Santa Cruz; sc-373759; 1:100 dilution or Abcam; ab21356; 1:100 dilution). Images were acquired using a laser-scanning confocal microscope (Zeiss LSM 880 or Olympus FV3000) as z-stack images and analyzed using ImageJ software (NIH).

### Statistical Analysis

Data are presented as means ± SD. Statistical tests were performed using Prism software, with methods described in the Figure Legends.

## Acknowledgments

We thank F. Tsuruta (Tsukuba University) and T. Masuda (Kyushu University) for critical reading; The University of Tokyo IQB Olympus Bioimaging Center (TOBIC) for microscopy; E. Ogawara (The University of Tokyo) for technical assistance; and members of Gotoh and Kishi laboratories for helpful discussion.

## Funding

This research was supported by AMED-CREST (JP23gm1310004 to Y.G.), AMED-PRIME (JP22gm6110021 to Y.K.), MEXT/JSPS KAKENHI (JP22H00431 to Y.G.; 16H06279, JP22H04687, 23H04214, and 24H01227 to Y.K.), the Takeda Science Foundation, the Uehara Memorial Foundation, the Asahi Glass Foundation, the Chugai Foundation for Innovative Drug Discovery Science, the Astellas Foundation for Research on Metabolic Disorders, the Naito Foundation, the SECOM Science and Technology Foundation, the Ono Pharmaceutical Foundation for Oncology, Immunology, and Neurology, and the Kurata Grants by The Hitachi Global Foundation.

## Author Contributions

Conceptualization, A. S. and Y. K.; Data Curation, A. S., H. M., and Y. K.; Formal Analysis, A. S. and H. M.; Funding Acquisition, Y. K.; Investigation, A. S., H. M., and Y. K.; Methodology, A. S., H. M., M. B., and Y. K.; Project Administration, Y. K.; Supervision, Y. K. and Y. G.; Visualization, A. S., H. M., and Y. K.; Writing – Original Draft Preparation, A. S. and Y. K.; Writing – Review & Editing, A. S., H. M., M. B., Y. G., and Y. K.

## Data Availability

The datasets generated during the current study are available from the corresponding author on reasonable request. The IDs of public scRNA-seq and snRNA-seq data were provided in Supplementary Table 1.

## Declaration of Interests

The authors declare no competing interests.

---

**Extended Data Figure 1.**
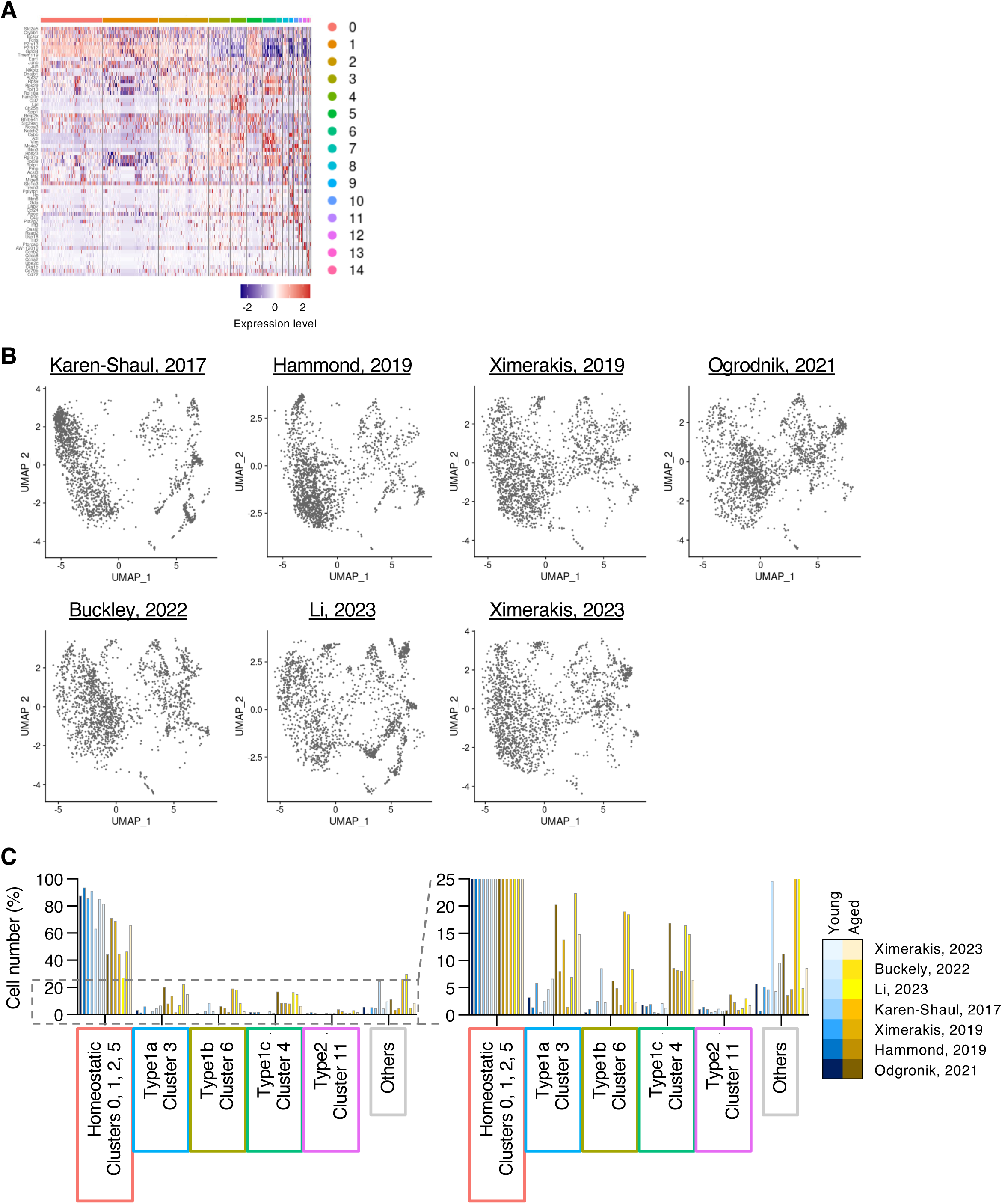
Integration of seven public scRNA-seq datasets of young and aged microglia, related to. **Figure 1**. **A.** Distribution of microglia from each dataset in UMAP space. **B.** Percentages of homeostatic microglia and common AAMs in each dataset.

**Extended Data Figure 2.**
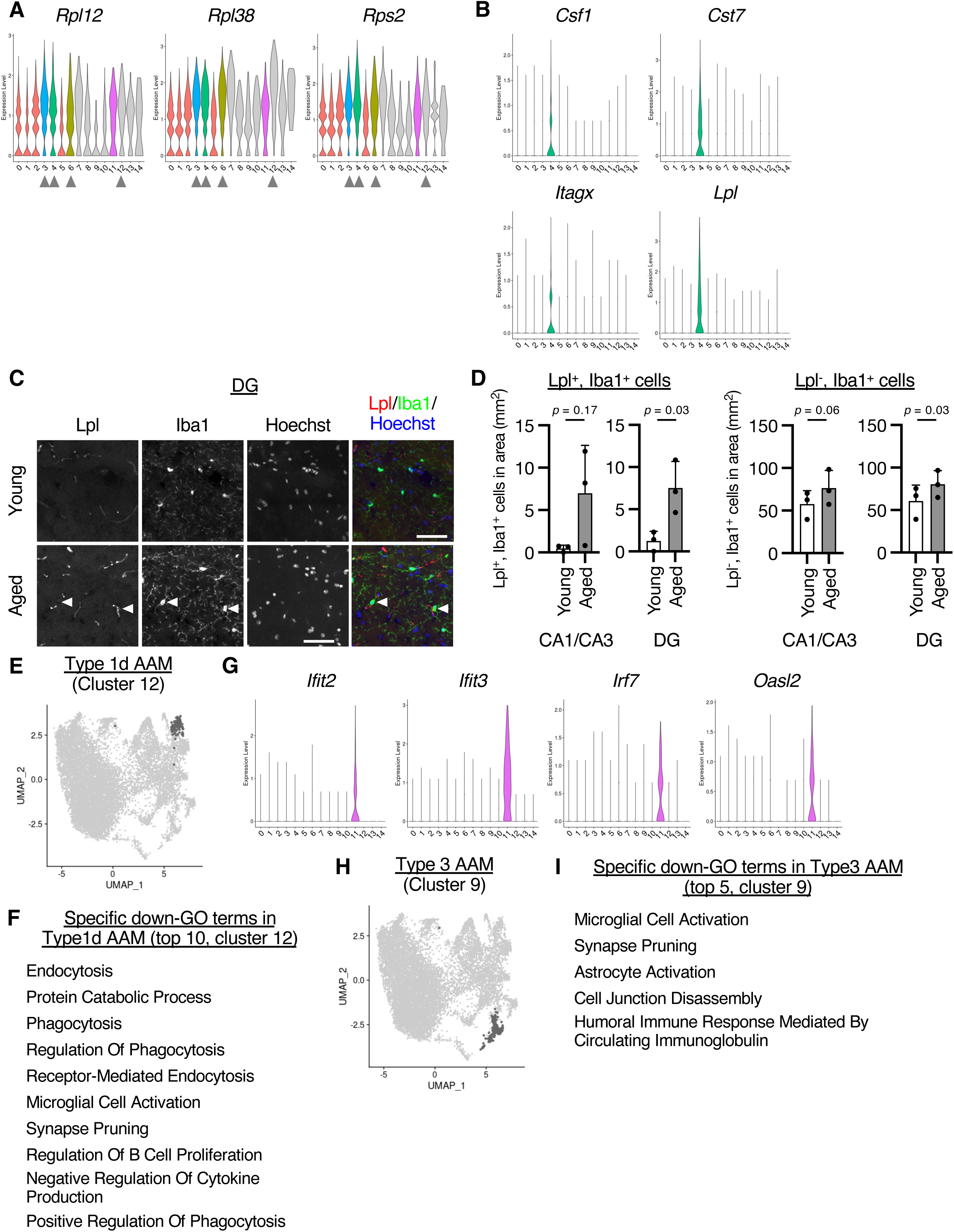
Characteristics of common and context-dependent AAMs, related to Figure 2. **A–B**. Distribution of microglia in the two context-dependent AAMs in UMAP space. **C.** Expression levels of ribosomal genes (*Rpl12*, *Rpl38*, and *Rps2*) in each cluster. **D.** Expression levels of upregulated DEGs in Type1c AAMs (*Csf1*, *Cst7*, Itgax, and *Lpl*) in each cluster. **E.** Coronal sections of young and aged brains stained with antibodies against Lpl and Iba1. Nuclei were counterstained with Hoechst 33342. Images were obtained from the dentate gyrus (DG). Arrowheads indicate Lpl+ and Iba1+ cells. Scale bars, 50 μm. **F.** Quantification of Lpl-positive (left panel) and Lpl-negative (right panel) Iba1-positive cells in the CA1/CA3 and DG regions. Data from three independent experiments are presented as mean + SD. *p*-values were calculated using paired Student’s t-test. **G.** Top ten Biological Process GO terms of upregulated DEGs in Type1d AAMs. **H.** Expression levels of upregulated DEGs in Type2 AAMs (*Ifit2*, *Ifit3*, *Irf7*, and *Oasl2*) in each cluster. **I.** Top five Biological Process GO terms of upregulated DEGs in Type2 AAMs.

**Extended Data Figure 3.**
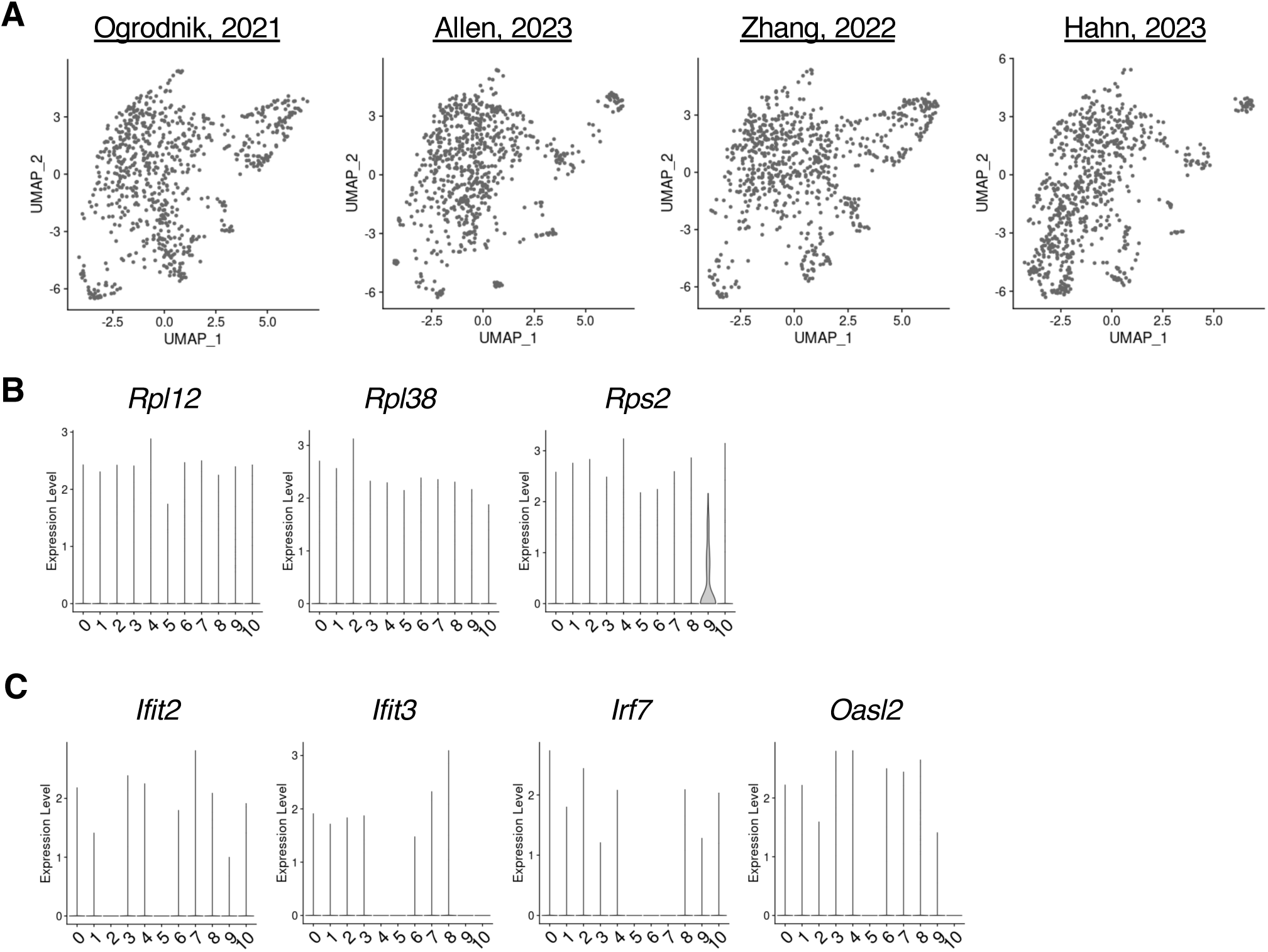
Integration of four public snRNA-seq datasets of young and aged microglia, related to. **Figure 3**. **A.** Distribution of microglia from each dataset in UMAP space. **B.** Expression levels of ribosomal genes (*Rpl12*, *Rpl38*, and *Rps2*) in each cluster. **C.** Expression levels of upregulated DEGs in Type2 AAMs (*Ifit2*, *Ifit3*, *Irf7*, and *Oasl2*) in each cluster.

**Extended Data Figure 4.**
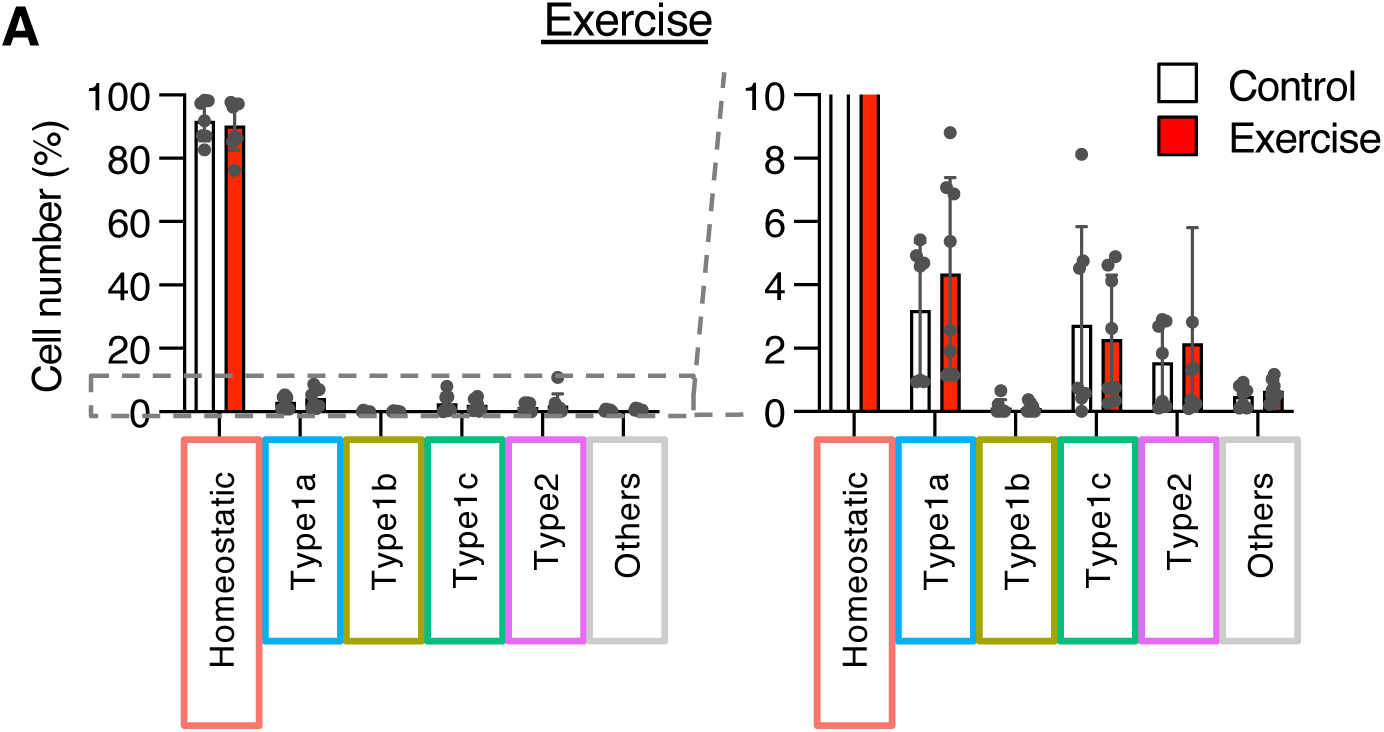
The effect of rejuvenation and premature aging manipulations on the number of homeostatic microglia and four common AAMs, related to. **Figure 6**. **A.** Prediction of corresponding clusters (homeostatic microglia, four common AAMs, and others) for cells from scRNA-seq of mice subjected to exercise (D). Bar graphs show the number of predicted cells. Data are presented as mean ± SD.

**Extended Data Figure 5.**
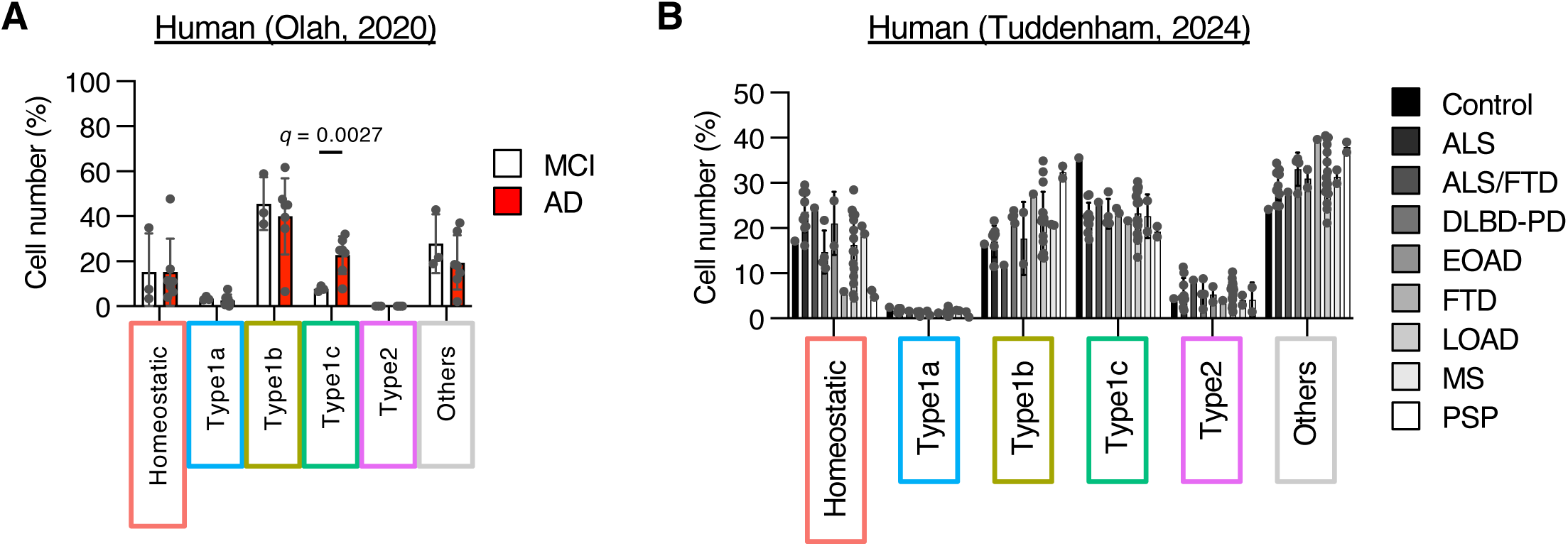
Corresponding microglia of mouse AAMs in human brains. **A–B**. Prediction of corresponding clusters (homeostatic microglia, four common AAMs, and others) for cells from human scRNA-seq datasets of neurological diseases. Bar graphs show the number of predicted cells. Data are presented as mean ± SD. p-values were calculated using multiple unpaired t-tests with Welch’s correction. MCI, mild cognitive impairment. AD, Alzheimer’s disease. ALS, amyotrophic lateral sclerosis. FTD, frontotemporal dementia. DLBP-PD, diffuse Lewy body disease-Parkinson’s disease. EOAD, early-onset AD. LOAD, late-onset AD. PSP, progressive supranuclear palsy.

## Notes

### Competing Interest Statement

The authors have declared no competing interest.

## References

1. Hou, Y. et al. Ageing as a risk factor for neurodegenerative disease. Nat Rev Neurol 15, 565–581 (2019).

2. Mosher, K. I. & Wyss-Coray, T. Microglial dysfunction in brain aging and Alzheimer’s disease. Biochem. Pharmacol. 88, 594–604 (2014).

3. Antignano, I., Liu, Y., Offermann, N. & Capasso, M. Aging microglia. Cell. Mol. Life Sci. 80, 126 (2023).

4. Ye, S.-M. & Johnson, R. W. Increased interleukin-6 expression by microglia from brain of aged mice. J. Neuroimmunol. 93, 139–148 (1999).

5. Sierra, A., Gottfried Blackmore, A. C., McEwen, B. S. & Bulloch, K. Microglia derived from aging mice exhibit an altered inflammatory profile. Glia 55, 412–424 (2007).

6. Godbout, J. P. et al. Exaggerated neuroinflammation and sickness behavior in aged mice after activation of the peripheral innate immune system. FASEB J. 19, 1329–1331 (2005).

7. Nakanishi, H. & Wu, Z. Microglia-aging: Roles of microglial lysosome- and mitochondria-derived reactive oxygen species in brain aging. Behav. Brain Res. 201, 1–7 (2009).

8. Marschallinger, J. et al. Lipid-droplet-accumulating microglia represent a dysfunctional and proinflammatory state in the aging brain. Nat Neurosci 23, 194–208 (2020).

9. Njie eMalick G., et al. Ex vivo cultures of microglia from young and aged rodent brain reveal age-related changes in microglial function. Neurobiol. Aging 33, 195.e1–195.e12 (2012).

10. Floden, A. M. & Combs, C. K. Microglia Demonstrate Age-Dependent Interaction with Amyloid-β Fibrils. J. Alzheimer’s Dis. 25, 279–293 (2011).

11. Gabandé Rodríguez, E., Keane, L. & Capasso, M. Microglial phagocytosis in aging and Alzheimer’s disease. J. Neurosci. Res. 98, 284–298 (2020).

12. Hammond, T. R. et al. Single-Cell RNA Sequencing of Microglia throughout the Mouse Lifespan and in the Injured Brain Reveals Complex Cell-State Changes. Immunity 50, 253–271.e6 (2019).

13. Frigerio, C. S. et al. The Major Risk Factors for Alzheimer’s Disease: Age, Sex, and Genes Modulate the Microglia Response to Aβ Plaques. Cell Rep. 27, 1293–1306.e6 (2019).

14. Safaiyan, S. et al. White matter aging drives microglial diversity. Neuron 109, 1100–1117.e10 (2021).

15. Kaya, T. et al. CD8+ T cells induce interferon-responsive oligodendrocytes and microglia in white matter aging. Nat. Neurosci. 25, 1446–1457 (2022).

16. Allen, W. E., Blosser, T. R., Sullivan, Z. A., Dulac, C. & Zhuang, X. Molecular and spatial signatures of mouse brain aging at single-cell resolution. Cell 186, 194–208.e18 (2023).

17. Hahn, O. et al. Atlas of the aging mouse brain reveals white matter as vulnerable foci. Cell 186, 4117–4133.e22 (2023).

18. Keren-Shaul, H. et al. A Unique Microglia Type Associated with Restricting Development of Alzheimer’s Disease. Cell 169, 1276–1290.e17 (2017).

19. Zhang, W., Sun, H.-S., Wang, X., Dumont, A. S. & Liu, Q. Cellular senescence, DNA damage, and neuroinflammation in the aging brain. Trends Neurosci. 47, 461–474 (2024).

20. Villeda, S. A. et al. The ageing systemic milieu negatively regulates neurogenesis and cognitive function. Nature 477, 90–94 (2011).

21. Villeda, S. A. et al. Young blood reverses age-related impairments in cognitive function and synaptic plasticity in mice. Nature Medicine 20, 659–663 (2014).

22. Katsimpardi, L. et al. Vascular and Neurogenic Rejuvenation of the Aging Mouse Brain by Young Systemic Factors. Science 344, 630–634 (2014).

23. Mahmoudi, S., Xu, L. & Brunet, A. Turning back time with emerging rejuvenation strategies. Nat. Cell Biol. 21, 32–43 (2019).

24. Wu, Y. E., Pan, L., Zuo, Y., Li, X. & Hong, W. Detecting Activated Cell Populations Using Single-Cell RNA-Seq. Neuron 96, 313–329.e6 (2017).

25. Ogrodnik, M. et al. Whole-body senescent cell clearance alleviates age-related brain inflammation and cognitive impairment in mice. Aging cell e13296 (2021) doi:10.1111/acel.13296.

26. Ximerakis, M. et al. Single-cell transcriptomic profiling of the aging mouse brain. Nature Neuroscience 22, 1696–1708 (2019).

27. Ximerakis, M. et al. Heterochronic parabiosis reprograms the mouse brain transcriptome by shifting aging signatures in multiple cell types. Nat Aging 3, 327–345 (2023).

28. Li, X. et al. Transcriptional and epigenetic decoding of the microglial aging process. *Nat*. Aging 3, 1288–1311 (2023).

29. Buckley, M. T. et al. Cell-type-specific aging clocks to quantify aging and rejuvenation in neurogenic regions of the brain. Nat Aging 3, 121–137 (2023).

30. Stuart, T. et al. Comprehensive Integration of Single-Cell Data. Cell 177, 1888–1902.e21 (2019).

31. Holtman, I. R. et al. Induction of a common microglia gene expression signature by aging and neurodegenerative conditions: a co-expression meta-analysis. Acta Neuropathol. Commun. 3, 31 (2015).

32. Keane, L. et al. mTOR-dependent translation amplifies microglia priming in aging mice. J. Clin. Investig. 131, (2020).

33. Blumenfeld, J., Yip, O., Kim, M. J. & Huang, Y. Cell type-specific roles of APOE4 in Alzheimer disease. Nat. Rev. Neurosci. 25, 91–110 (2024).

34. Krasemann, S. et al. The TREM2-APOE Pathway Drives the Transcriptional Phenotype of Dysfunctional Microglia in Neurodegenerative Diseases. Immunity 47, 566–581.e9 (2017).

35. Zhang, Y. et al. Single-cell epigenome analysis reveals age-associated decay of heterochromatin domains in excitatory neurons in the mouse brain. Cell Res 1–14 (2022) doi:10.1038/s41422-022-00719-6.

36. Thrupp, N. et al. Single-Nucleus RNA-Seq Is Not Suitable for Detection of Microglial Activation Genes in Humans. Cell Rep. 32, 108189 (2020).

37. Reddy, P. H. & Beal, M. F. Amyloid beta, mitochondrial dysfunction and synaptic damage: implications for cognitive decline in aging and Alzheimer’s disease. Trends Mol. Med. 14, 45–53 (2008).

38. Oakley, H. et al. Intraneuronal β-Amyloid Aggregates, Neurodegeneration, and Neuron Loss in Transgenic Mice with Five Familial Alzheimer’s Disease Mutations: Potential Factors in Amyloid Plaque Formation. J. Neurosci. 26, 10129–10140 (2006).

39. Beuker, C. et al. Stroke induces disease-specific myeloid cells in the brain parenchyma and pia. Nat. Commun. 13, 945 (2022).

40. Gulen, M. F. et al. cGAS–STING drives ageing-related inflammation and neurodegeneration. Nature 620, 374–380 (2023).

41. Olah, M. et al. Single cell RNA sequencing of human microglia uncovers a subset associated with Alzheimer’s disease. Nat. Commun. 11, 6129 (2020).

42. Tuddenham, J. F. et al. A cross-disease resource of living human microglia identifies disease-enriched subsets and tool compounds recapitulating microglial states. Nat. Neurosci. 27, 2521–2537 (2024).

43. Silvin, A. et al. Dual ontogeny of disease-associated microglia and disease inflammatory macrophages in aging and neurodegeneration. Immunity 55, 1448–1465.e6 (2022).

44. Harrison, D. E. et al. Rapamycin fed late in life extends lifespan in genetically heterogeneous mice. Nature 460, 392–395 (2009).

45. Neff, F. et al. Rapamycin extends murine lifespan but has limited effects on aging. J. Clin. Investig. 123, 3272–3291 (2013).

46. Malagelada, C., Jin, Z. H., Jackson-Lewis, V., Przedborski, S. & Greene, L. A. Rapamycin Protects against Neuron Death in In Vitro andIn Vivo Models of Parkinson’s Disease. J. Neurosci. 30, 1166–1175 (2010).

47. Spilman, P. et al. Inhibition of mTOR by Rapamycin Abolishes Cognitive Deficits and Reduces Amyloid-β Levels in a Mouse Model of Alzheimer’s Disease. PLoS ONE 5, e9979 (2010).

48. Masuda, T. et al. Spatial and temporal heterogeneity of mouse and human microglia at single-cell resolution. Nature 566, 388–392 (2019).

49. Chen, E. Y. et al. Enrichr: interactive and collaborative HTML5 gene list enrichment analysis tool. BMC Bioinform. 14, 128 (2013).

50. Frey, T. et al. Age[associated reduction of nuclear shape dynamics in excitatory neurons of the visual cortex. Aging Cell e13925 (2023) doi:10.1111/acel.13925.

